# Genetics of intra-species variation in avoidance behavior induced by a thermal stimulus in *C. elegans*

**DOI:** 10.1101/014290

**Authors:** Rajarshi Ghosh, Joshua S. Bloom, Aylia Mohammadi, Molly E. Schumer, Peter Andolfatto, William Ryu, Leonid Kruglyak

## Abstract

Individuals within a species vary in their responses to a wide range of stimuli, partly as a result of differences in their genetic makeup. Relatively little is known about the genetic and neuronal mechanisms contributing to diversity of behavior in natural populations. By studying animal-to-animal variation in innate avoidance behavior to thermal stimuli in the nematode *Caenorhabditis elegans*, we uncovered genetic principles of how different components of a behavioral response can be altered in nature to generate behavioral diversity. Using a thermal pulse assay, we uncovered heritable variation in responses to a transient temperature increase. Quantitative trait locus mapping revealed that separate components of this response were controlled by distinct genomic loci. The loci we identified contributed to variation in components of thermal pulse avoidance behavior in an additive fashion. Our results show that the escape behavior induced by thermal stimuli is composed of simpler behavioral components that are influenced by at least six distinct genetic loci. The loci that decouple components of the escape behavior reveal a genetic system that allows independent modification of behavioral parameters. Our work sets the foundation for future studies of evolution of innate behaviors at the molecular and neuronal level.

## Introduction

Individuals vary widely in their behavioral responses to a given stimulus. Differences in environment, the genetic variation between individuals or both are thought to contribute to variation in behavior. Though significant progress has been made in understanding the genetic basis of behavior using techniques such as inducing mutations in a single genetic background [1-6], much less is known about the genetic basis of behavioral variation in natural populations.

One challenge in understanding the genetic basis of behavior is comprehensively capturing the behavior of interest. Genetic studies of behavior generally focus on a single summary phenotypic metric. However, several distinct motor outputs are integrated over time to give rise to a behavioral response to a given stimulus [7-11]. With high resolution phenotyping behavioral responses to a particular stimulus can be broken down into simpler components or sub-behaviors. Diversity in the behavioral responses to a given stimulus in natural populations could result from modifications of the sub-behaviors through natural selection, as postulated by Darwin ~200 years ago [12]. By quantitatively describing sub-behaviors and their variation within and between species, Tinbergen and others were able to identify ample heritable variation in sub-behaviors in multiple species [13]. These studies laid the framework for understanding the origin and divergence of behaviors within and between species. However, until recently, we did not have the tools to ask questions about the genetic basis of sub-behaviors and the mechanisms through which they are integrated to generate behavioral variation.

The nematode *C. elegans* is the only organism whose complete neuronal connectivity is known. With well-established genetic and genomic tools and the availability of a large densely genotyped collection of natural isolates [14], *C. elegans* provides a unique opportunity to study the genetic basis of animal to animal variation in behavior. Tools for dissecting the operation of neuronal circuitry in this organism are also well developed [15,16], thus providing the ideal platform for connecting diversity at the genetic and neuronal circuit level to variation in behavior.

Here, we asked how genetic variation within a population contributes to differences in innate behavioral patterns. Specifically, we focused on the escape response of *C. elegans* to a noxious thermal stimulus, a quintessential innate behavior. Upon experiencing noxious thermal stimuli, *C. elegans* animals display a stereotypical behavioral sequence that can be readily broken down into several distinct behavioral components [1,7]. By combining well developed genetic and genomic resources of *C. elegans* with a thermal pulse assay [17,18], we characterized the genetic basis of natural variation in avoidance responses to thermal stimuli.

We used high-content behavioral phenotyping to capture the avoidance behavior of the global *C. elegans* population in response to thermal stimuli. Instead of considering a single metric summarizing the avoidance response, we simultaneously quantified multiple aspects of the avoidance response of animals exposed to thermal pulse stimuli. Additionally, we used linkage analysis to identify distinct genetic loci underlying different aspects of the escape behavioral pattern. Our results, consistent with observations in other organisms [19,20], illustrate that genetically separate modules allow for individual variation in sub-behaviors that together result in behavioral diversity within the *C. elegans* species. Our results provide novel insights into the genetic mechanisms regulating behavior, the operation of the nervous system and the evolution of avoidance behaviors.

## Results

### Wild isolates of *C. elegans* differ in multiple aspects of escape behavior induced by thermal pulse stimulus

Upon sensation of a noxious thermal stimulus, *C. elegans* animals typically move backwards, turn, and resume forward movement in a new direction (Fig. 1A,Video S1-4). We characterized multiple aspects of the avoidance behavior in response to a thermal pulse that transiently raises the temperature surrounding a forward moving animal [7]. Briefly, we recorded the behavior of each animal for 15 seconds in response to thermal pulse stimuli of various intensities (Fig. 1A) and quantified sixty-six metrics that characterized the resulting escape behavior (see Methods) for each thermal pulse stimulus. To investigate the genetic basis of individual variation in response to thermal stimuli, we first surveyed the thermal pulse induced behavioral responses of 89 strains of *C. elegans* isolated from different geographical regions across the globe (Fig. 1B) [14]. These animals exhibited heritable variation in multiple aspects of their escape behavioral pattern in response to a thermal pulse of 0.4°C above the baseline temperature (ΔT = 0.4°C) (Fig. 1B). Depending on the sub-behavior analyzed, the broad-sense heritability among the wild isolate population ranged from 0.04 to 0.21 (Figure S5A). The heritability is in a range consistent with that observed for other behavioral responses to a noxious, stressful stimulus [21-23]. We found weak correlations among certain escape behavior components in the wild isolate populations, suggesting a separate genetic basis for these traits (Figure S5B). For example, the speed at an intermediate time point (1.61 second) was only weakly correlated (r=0.242) with the speed at an early time point (0.18 second) in the wild isolate population. Conversely, the speeds at early (0.18 second) and late (8.1 second) time points were strongly correlated (r=0.874) (Figure S5B).

**Figure 1:**
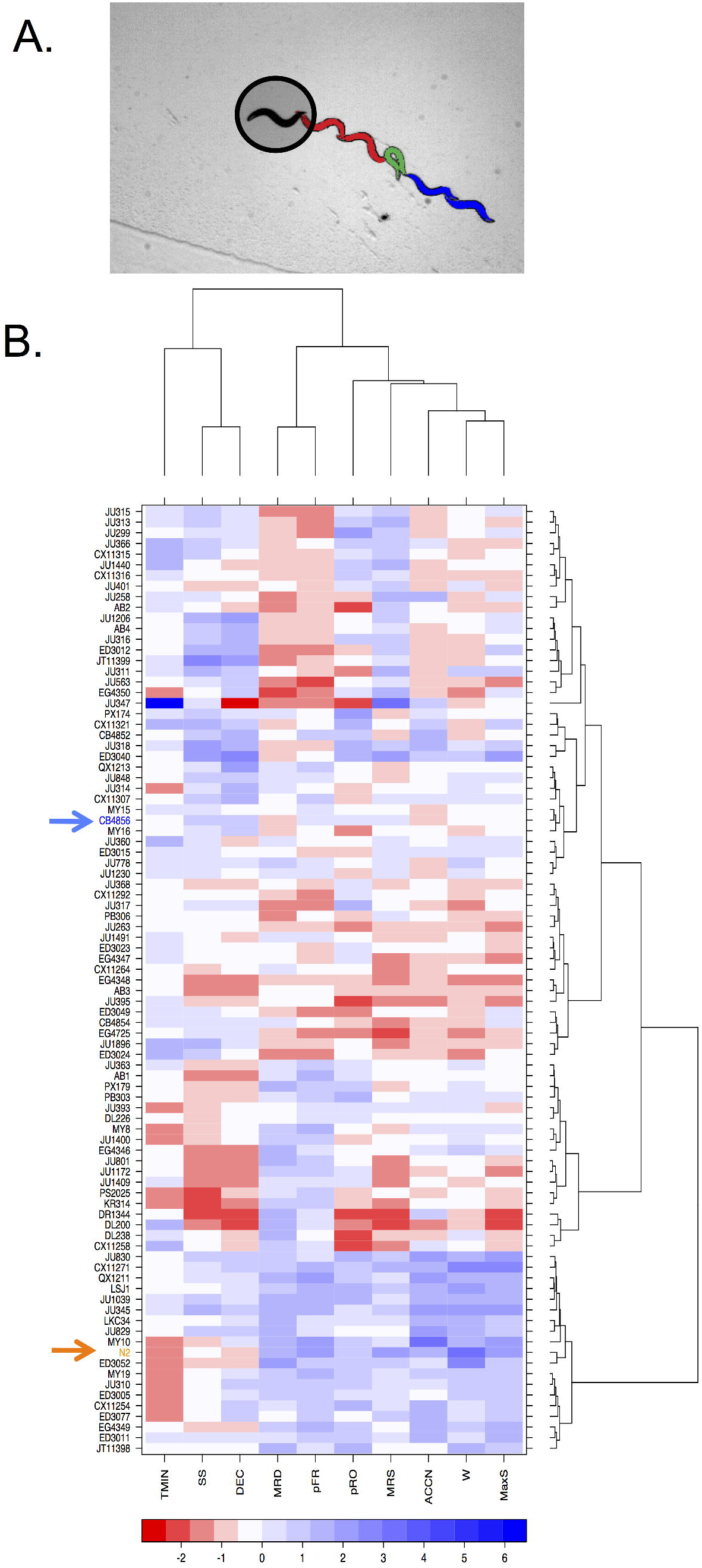
Variation in components of avoidance behavior among wild isolates of *C. elegans* in response to thermal pulse. A) The thermal avoidance assay and response. An infrared laser with a beam width larger than the worm body heats the surrounding area of the animal. The typical responses constitute a reversal (red), an omega turn (green), and forward movement (blue). B) Profiles of behavioral metrics of 89 wild isolates of *C. elegans* in response to ΔT~0.4°C. Each behavioral feature is Z-score normalized (See Methods). TMIN: time to reach pause state, SS: speed at start of the thermal pulse, DEC: deceleration to the pause state, MRD: mean reversal duration, pFR: probability of responding by reversal, pRO: probability of responding by omega, MRS: mean reversal speed, ACCN: acceleration, W: number of body bends during reversals, MaxS: Peak speed after exposure to thermal pulse. In the heatmaps, the strains are reordered by the dendrograms resulting from hierarchical clustering. The Bristol strain (N2) and the Hawaiian strain (CB4856) are depicted by arrows and are colored orange and blue respectively. For simplicity we show only 10 behavioral components. The other 56 behavioral metrics from the centroid speed profiles are not shown.

To determine the genetic architecture underlying diversity in escape behavior in global populations of *C. elegans*, we conducted genome-wide association mapping of the sixty-six behavioral metrics that describe the escape response to a thermal stimulus of 0.4°C. For the majority of traits, we were unable to detect significant associations. However, loci on chromosomes IV and X contributed to the variation in acceleration, whereas loci on chromosome X contributed to variation in pre-stimulus speed (Figure S5C-D), suggesting that these two traits may be genetically separable.

### Animals from Hawaii and Bristol differ in their responses to ΔT = 0.4°C

Among the strains analyzed for genome-wide association we found that relative to a strain from Hawaii, the laboratory strain (originally from Bristol) differed significantly in multiple aspects of the escape behavioral pattern in response to ΔT = 0.4°C (Fig. 1B). These two strains exhibited distinct avoidance response behavioral profiles (Fig. 1B, S1-4).

In agreement with previous studies [7,17], we found that in un-stimulated conditions and immediately (< 0.2 seconds) after exposure to thermal stimuli, Bristol animals move with a significantly lower speed compared to the Hawaiian animals (Fig. 2A, top panel). Upon experiencing the thermal pulse, both strains decelerate to a pause state and then accelerate away from the stimulus. However, the two strains showed remarkably different speed profiles during the avoidance response (Fig. 2A). Bristol and Hawaiian strains also differed in their probability of transition between different behavioral states, as well as in the duration of a given behavioral state (Fig. 2B). At stimuli with higher intensities (ΔT of 1°C, 4.8°C or 9.1°C), the differences in avoidance response between these two strains were smaller (Fig. 2B, S1-4). This result suggests that the differences in behavioral metrics we observed at ΔT = 0.4°C are not a reflection of differences in general locomotion between these two strains. Because the parental strains differed most in their responses elicited by ΔT = 0.4°C, we focused our studies of the genetic basis in avoidance behavior at this stimulus intensity.

**Figure 2:**
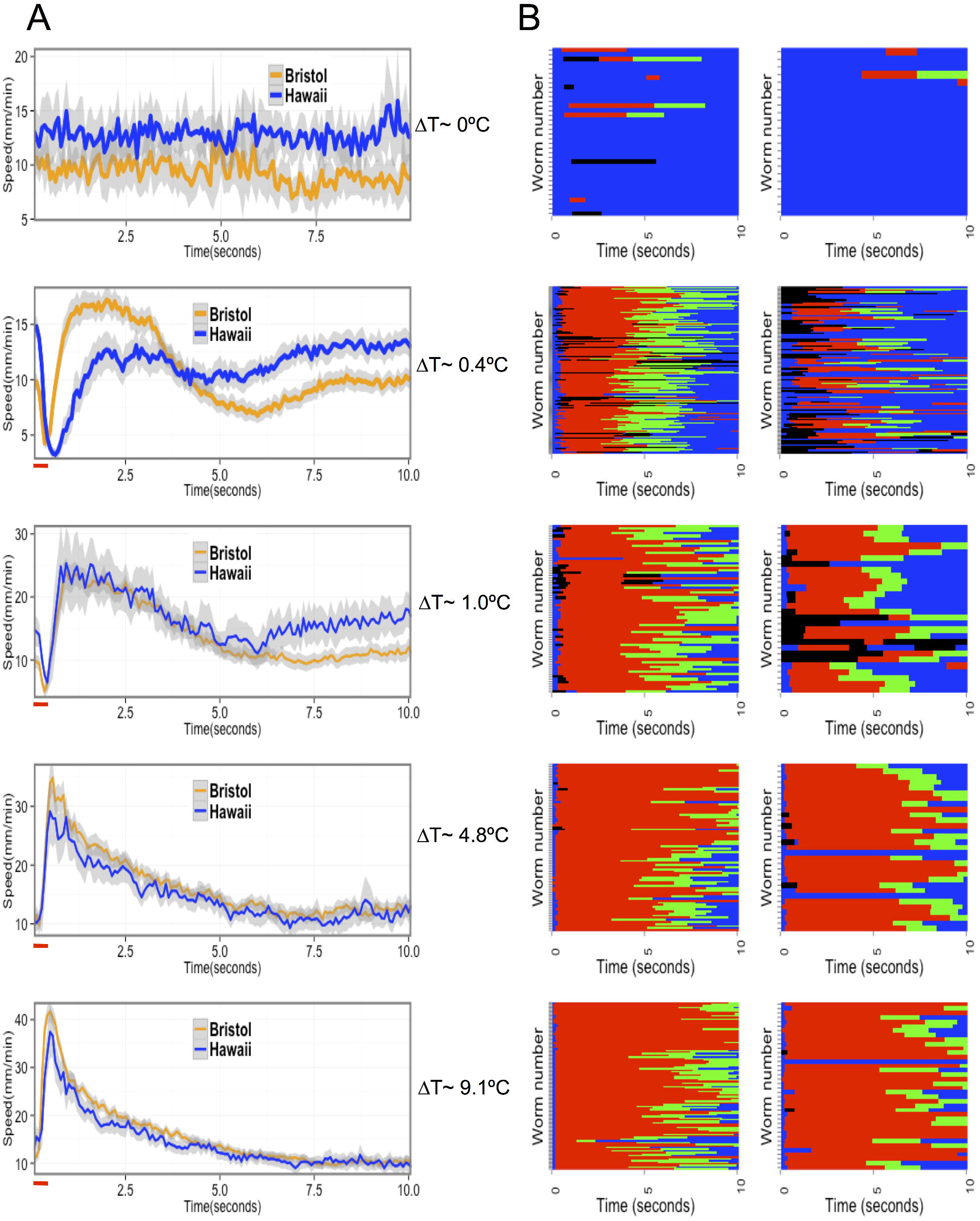
Comparison of Hawaii and Bristol strains in responses to thermal stimuli of various intensities. A) Centroid speed of Bristol (orange) and Hawaii (blue) animals plotted against time for each ΔT. The red horizontal bar indicates duration of the pulse. Temperature increases above the baseline (ΔT) as a result of thermal pulses are indicated. The resulting speed profile is in the panel corresponding to the indicated ΔT. (B) Ethogram of different behavioral states of the Bristol (left) and Hawaii (right) animals at indicated ΔT. The behavioral sequence of each animal over the duration of the assay at a given ΔT is shown. Each row represents behavior of a single animal over time. Blue = forward state, red = reversal, black = pause, green = omega turns.

### Behavioral differences between Hawaii and Bristol strains are heritable

We characterized the individual components of the escape response of 138 Recombinant inbred lines (RIAILs), made from Bristol and Hawaii parent strains [24]. We found large phenotypic variation in different sub-behaviors among the RIAILs. Broad-sense heritability ranged from 0.04 to 0.27, similar to estimates obtained from the wild isolates for the same set of behavioral metrics (Figure S5A). To identify loci underlying these traits, we conducted QTL mapping with behavioral metrics extracted from the centroid speed profile (Fig. 2A, second panel from top, see Methods) and the ethograms (Fig. 2B second panel from top, see Methods).

### Distinct loci contribute to variation in speed profiles

The Bristol and Hawaiian strains differed significantly in their speed profiles in responses to ΔT = 0.4°C. Initially (<0.5 seconds from application of the thermal pulse), the Hawaiian animals moved forward at a greater speed compared to the Bristol animals, similar to observations under unstimulated conditions (Fig. 3A, top panel). Subsequently, the Bristol animals moved away from the stimulus significantly faster than the Hawaiian animals. After approximately three seconds, the speed differences between these two strains were similar to the pre-stimulus state, with the Hawaiian animals once again moving faster (Fig. 3A, upper panel). We used speed at 0.18 second increments for 10 seconds after the stimulus as a trait in linkage analysis and identified two loci, on chromosomes IV and X, that contributed to variation in the centroid speed profile among the RIAILs (Fig. 3A, Fig. S6 A-F).

**Figure 3:**
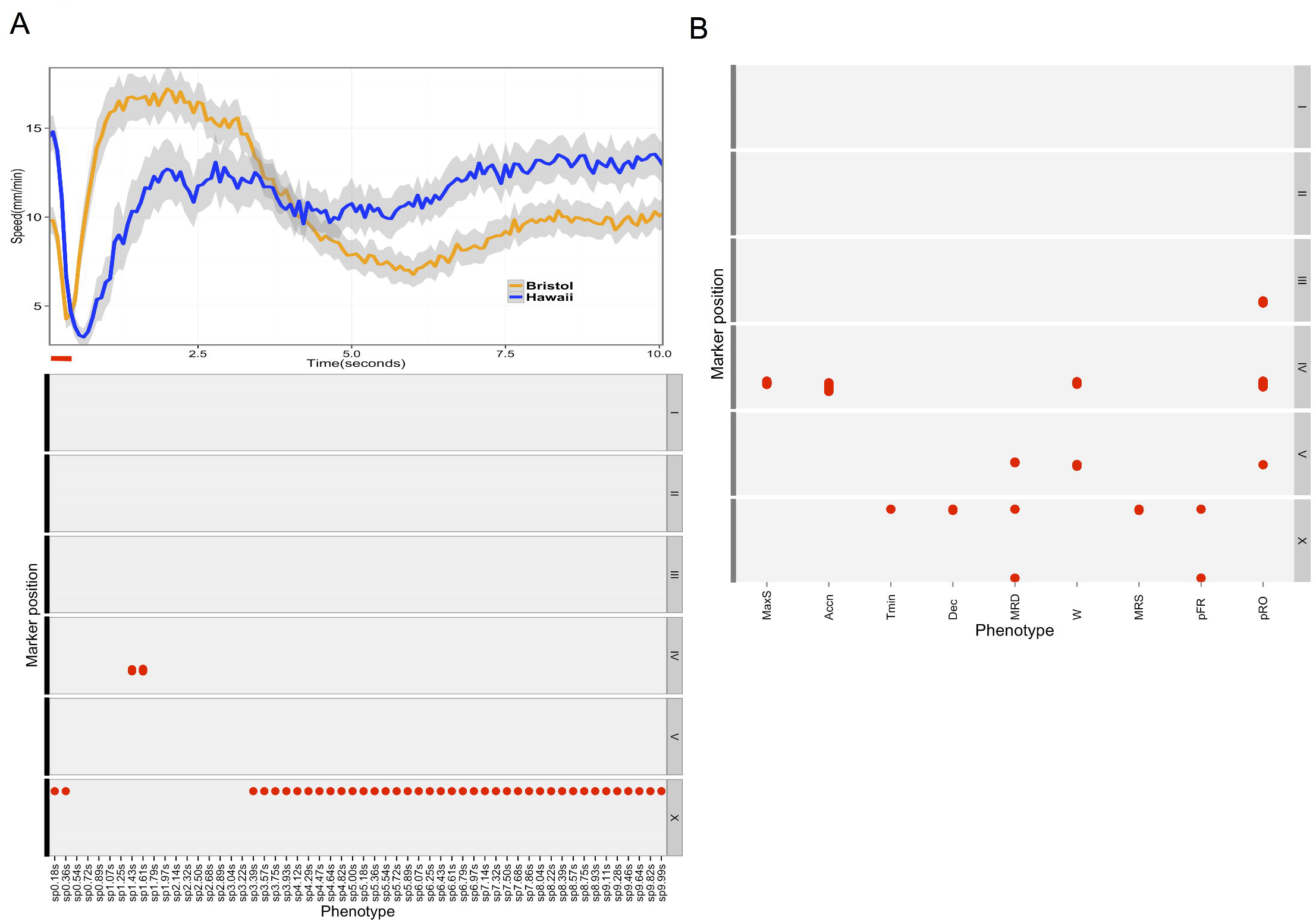
Genomic intervals contributing to variation in components of escape behavior in 138 RIAILs. A) Upper panel: Speed profile of Bristol (orange) and Hawaii (blue) strain (same as Fig 2A, panel 2). The thermal pulse is depicted by a red horizontal line. Lower panel: The red dot represents position of significant QTL associated with speed at every 0.18 seconds (X-axis) at the corresponding marker position plotted on the Y axis. X axis depicts the traits (the speed profile treated as behavioral metrics) used for linkage mapping. sp*.**s represents speed at *.** seconds from start of the assay. Y axis: Marker position for each chromosome. The chromosomes I though X are shown on the left of each facet. B) The red dots represent position of significant QTL associated with speed at every 0.18 seconds (X-axis) at the corresponding marker position plotted on the Y axis (same as A). X axis depicts the traits used for linkage mapping with 138 RIAILs.

A single significant QTL on chromosome X contributed to variation in speeds at early (< 0.5 seconds from the thermal pulse) and late (> 3.39 seconds from the thermal pulse application) time points (Fig 3A, Fig. S6 A,C,D,F). This locus explained approximately 47% and 33% of variation in average speeds at 0.18 and 5.0 seconds, respectively. The 1.5-LOD support interval of this QTL includes the gene *npr-1*; polymorphism in this gene has been shown to underlie speed differences between the Bristol and Hawaiian strains. We also identified a QTL on chromosome IV that contributed to variation in speed at 1.43 and 1.61 seconds after the thermal pulse (Fig. 3A, Fig. S6 B, E). This locus explained ~12% of the variance in speed. We did not detect a significant QTL on chromosome X for speeds at 1.43 and 1.61 seconds (Fig. 3A, Compare Fig. S6 E,F), suggesting that the two loci contribute to variation in speed at different phases of the response.

An alternative way to describe the escape behavior is to quantify discrete aspects of the speed profile. Upon exposure to a thermal pulse, forward moving animals decelerate to a pause state, after which they accelerate away from the stimulus (Fig. 3A, upper panel). Therefore, the centroid speed profile can be divided into an initial deceleration period followed by an acceleration phase. From the speed profiles, we quantified the maximum speed reached after exposure to a thermal pulse, as well as acceleration to maximum speed, defined as the rate of change in the centroid speed from the pause state to the maximum speed. For peak speed and acceleration, we identified a QTL on chromosome IV, at the same location as obtained with speeds at 1.43 and 1.61 seconds (Fig. 3B, Fig. S6 G,H). This locus explained ~13% of the variation in acceleration and peak speed among the RIAILs. We also quantified the time it takes to reach the pause state as well as deceleration, defined as the rate of change in the centroid speed from the speed at the start of the assay to the pause state. We found that, in contrast to acceleration, overlapping QTL on chromosome X contributed to deceleration and time taken to reach the pause state (Fig. 3B, Fig. S6 I, J). These loci explained 10.5% and 11.2% of the variation in deceleration and time to minimum speed, respectively, in the RIAIL population. Thus acceleration and deceleration phases during avoidance response behavior are under independent genetic control.

Scanning for additional QTL with markers at the peaks of the two identified QTL used as covariates did not reveal any additional loci. Additionally, to scan for interactions between identified QTLs and other regions of the genome in controlling escape behavior, we performed a 2-D genome scan. We did not find evidence for any significant QTL-QTL interactions. Taken together, these results suggest that polymorphisms at two loci, on chromosomes IV and X, contribute additively to the observed variation in speed during thermal avoidance behavior in the RIAILs.

### Distinct loci contribute to variation in different avoidance behavior components

Several aspects of the avoidance behavior could not be fully captured by considering only the centroid of the animal alone. Upon sensation of a thermal stimulus, the posture of an animal changes drastically (Fig. 1A) during the reversals and omega turns. We obtained metrics that describe the reversals as well as the transition probabilities among forward, reversal and omega states (see Methods). We conducted QTL mapping separately for each of these traits and identified unique combinations of five distinct loci contributing to variation in these traits (Fig. 3B, Fig. S7 A-E).

We detected two distinct loci on chromosome X (QTL_Xa, QTL_Xb) that contributed to variation in the probability of responding by reversal (Fig. 3B, Fig S7A) and the average duration of reversals (Fig. 3B, Fig S7B), two traits that differ between the Hawaii and Bristol strains. For the average duration of reversals, we detected an additional QTL on chromosome V (QTL_V) (Fig. 3B, Fig. S7B) that did not significantly contribute to variation in the probability of responding by reversal (Fig 3B, Fig. S7A).

Two features of the reversal behavior—reversal duration and the number of body bends during reversal—were genetically separable. While the former was significantly influenced by QTL_Xa, QTL_Xb and QTL_V, variation in the latter was attributable to QTL on chromosome IV (QTL_IV) and QTL_V (Fig 3B, Fig S7C). QTL_IV contributed to variation in the number of body bends during reversal but not to the duration of reversals, suggesting that the underlying polymorphisms can alter the number of body bends without significantly changing reversal duration. Additionally, we identified QTL on chromosomes III, IV and V as significant contributors to variation in the probability of responding to the thermal stimulus with an omega turn. The interval of QTL on chromosomes IV and V for the probability of omega turns overlapped with QTL identified for other behavioral metrics (QTL_IV, QTL_V). The QTL on chromosome III was detected only for the probability of responding by omega (Fig 3B, Fig. S7D).

We detected QTL at overlapping positions using distinct behavioral measures. For example, QTL_IV, which affected the average number of body bends during reversals and the probability of responding with an omega turn, coincided with the QTL for peak speed and acceleration (Fig. 3A and B). Additionally, we found a significant QTL on chromosome X contributing to mean reversal speed (rate of propagation of body bends during reversal) (Fig. 3A,B, Fig. S7E). This QTL co-localized with the QTL on chromosome X detected for speeds at early and late time points (Fig. 3A). Using the peak marker on chromosome X as a covariate for reversal speed, we identified a QTL on chromosome IV (QTL_IV) at the same position as the QTL for peak speed, acceleration, and speeds at 1.43 and 1.61 seconds. This result is not unexpected, as the reversal speed calculated by dividing the number of body bends during reversal by reversal duration includes the acceleration phase of the escape response. We also detected QTL_IV for the number of body bends during reversals (Fig. 3B). Thus, using distinct behavioral measures, we detected the same loci.

We identified some loci (e.g. QTL_IV) affecting multiple behavioral parameters, whereas others (e.g. QTL_III) affected a single behavioral component of the escape response. Taken together, these data suggest that a modular genetic architecture contributes to diversity in thermal pulse escape response in *C. elegans*. This leads to the prediction that distinct combinations of the detected QTL will result in predictable changes in multiple behavioral metrics that together constitute an avoidance behavioral pattern. We next tested this behavioral prediction.

### Modular genetic architecture contributes to variation in behavioral responses to a thermal pulse

We identified two loci, QTL_IV and QTL_X, contributing to variation in behavioral components quantified from the centroid speed profile (Fig. 3A, Fig. S6). We next tested whether these two loci were sufficient to generate diversity in speed profiles during avoidance behavior, irrespective of the genetic background. We obtained the average speed profiles of a group of RIAILs that had Bristol alleles at the peak markers at QTL_IV and QTL_X. The speed profile of this group of RIAILs resembled the Bristol parental strain (Fig. 4A, red line, compare with Fig 3A, orange line in upper panel). Similarly, RIAILs with Hawaii alleles at QTL_X and QTL_IV exhibited speed profiles similar to the Hawaii parent (Fig. 4A, purple line, compare with blue line in Fig. 3A upper panel). QTL_IV contributes to variation in speed at intermediate time points, whereas QTL_X contributes at early and late time points. Non-parental combinations of alleles generated centroid profiles expected from a modular genetic architecture for speed. As expected, RIAILs with the Bristol allele of QTL_X and the Hawaii allele of QTL_IV exhibited speed profiles resembling the Bristol parent at early and late time points (Fig. 4A, green line) but the Hawaii parent at intermediate time points (Fig. 4A, green and purple lines). The reverse was true for RIAILs with the Hawaii allele of QTL_X and the Bristol allele of QTL_IV (Fig. 4A, cyan line). Thus the allelic states of QTL_IV and QTL_X define the shape of the centroid speed profile in a predictable manner. These results support the idea that a modular genetic architecture can generate individual variation in behavior, consistent with observations in other organisms [19,20].

**Figure 4:**
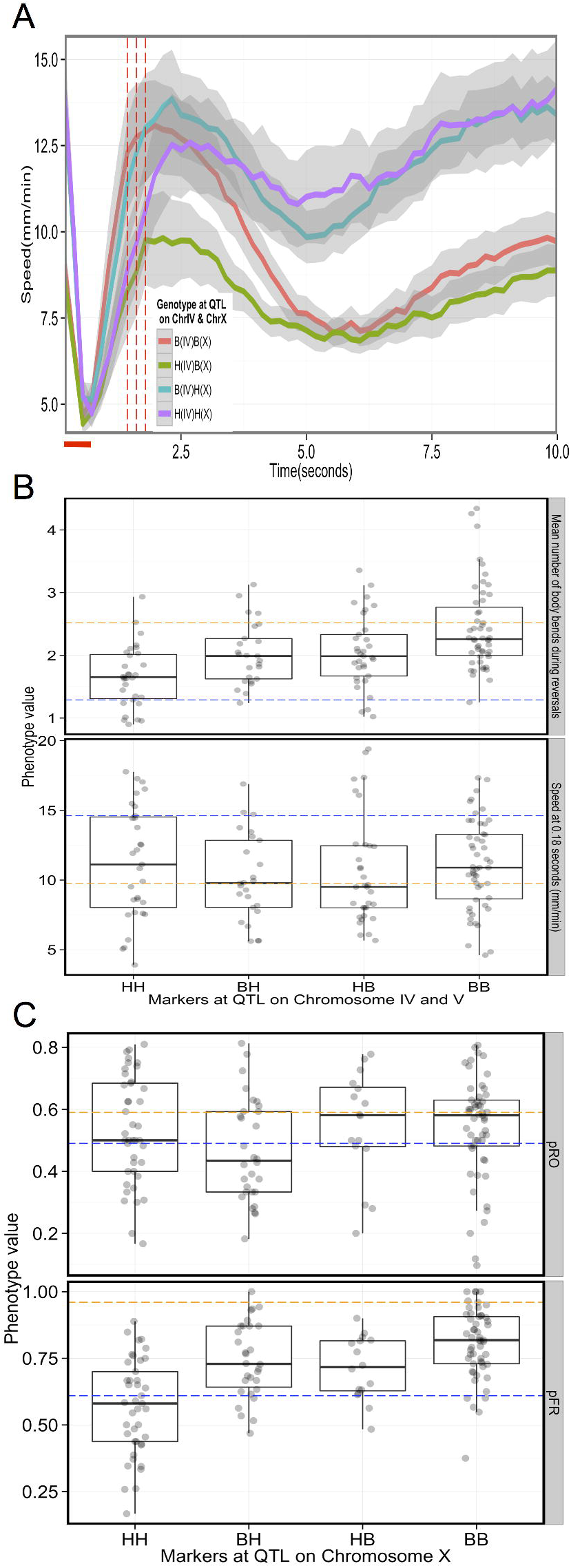
Modular genetic architecture contributes to variation in thermal avoidance behavior. A) Variation in speed profile is largely explained by two loci. The centroid speed plotted against time (seconds) of four groups of strains in response to ΔT~0.4°C. Strains with Bristol like alleles at the peak marker of the identified QTL on chromosome IV and X are depicted as B (IV) and B(X) whereas strains with Hawaii like alleles at the same markers are depicted as H(IV) and H(X). The three red dotted lines are the time points, speeds at which revealed a QTL on chromosome IV. The shaded area (gray) around each line represents the 95% confidence interval. The red horizontal line represents the duration of the laser pulse **B,C) Additive QTL contributes to variation in escape behavior** B)Mean number of body bends (upper panel) and speed at 0.18 seconds(lower panel) in groups of strains with Hawaii(HH) and Bristol(BB) alleles, Hawaii like allele at peak marker of QTL on chromosome IV and Bristol like allele at peak marker of QTL on chromosome IV(HB), Bristol like allele at peak marker of QTL on chromosome IV and Hawaii like allele at peak marker of QTL on chromosome IV(BH) at peak of detected QTL on chromosomes IV and V for these traits. The blue and orange dotted line represents the parental Hawaii and Bristol average phenotypes, respectively. C) Probability of omega turns (upper panel, pRO) and probability of responding by reversals (pFR, lower panel) in groups of strains with Hawaii (HH) and Bristol(BB) like alleles at peaks of the two QTL detected on chromosome X. BH: Bristol alleles at distal and Hawaii allele at the proximal QTL peak on chromosome X, HB: Hawaii alleles at distal and Bristol allele at the proximal QTL peak on chromosome X. The blue and orange dotted line represents the parental Hawaii and Bristol average phenotypes, respectively.

### QTL contribute additively to variation in escape behavior

We observed no evidence of epistasis for the suite of escape behavior components. The number of body bends during reversals of RIAILs with both Hawaii or both Bristol alleles at QTL_IV and QTL_V resembled that of Hawaii or Bristol parents, respectively (Fig. 4B, upper panel). RIAILs with Hawaii allele at QTL_IV and Bristol allele on QTL_V and the alternate allelic combination had intermediate phenotypes.

We detected two additive QTL on chromosome X that contributed to behavioral variation in the probability of responding by reversal, with the proximal QTL (QTL_Xa) explaining ~18% and the distal QTL (QTL_Xb) explaining ~7% of the phenotypic variation among the RIAILs. We observed an average probability of reversal of 0.56 (±0.02,N=41) in RIAILS with Hawaii alleles at these two loci on chromosome X (QTL_Xa and QTL_Xb), whereas the probability of responding by reversals for strains with Bristol alleles at these loci was 0.80 (±0.13,N=53), similar to the parental values. Strains with Hawaii alleles at QTL_Xa and Bristol alleles at QTL_Xb or the alternate allelic combination responded with an intermediate probability of reversals (Fig. 4C, lower panel). Taken together, our results suggest that distinct allelic combinations, acting in a largely additive fashion, contribute to variation in escape behavioral metrics.

### Fine mapping QTL using nearly isogenic lines

Variation in many avoidance behavior traits was controlled by a locus on chromosome X. This genomic region contributed to variation in speeds at early and late time points, deceleration, time to reach pause state, probability of responding by reversal and duration of reversals. A single pleiotropic gene or several linked functionally related genes could cause these phenotypic effects. To distinguish between these possibilities, we sought to identify the causative gene(s) at the QTL identified on chromosome X for these traits.

QTL contributing to variation in speed contained *npr-1*, a gene encoding a G-protein coupled receptor. A laboratory-derived gain-of-function variant of *npr-1* in the Bristol strain background has been previously shown to contribute to variation in several phenotypes [25-31], including a slower foraging speed relative to the Hawaii strain in the presence of food [25]. We found that F1 animals, obtained by reciprocally crossing the Bristol and Hawaii strains, exhibited speed at the start of the thermal pulse assay resembling Bristol animals, consistent with molecular gain of function in the Bristol strain underlying the observed speed differences (Fig. 5A). Consistent with this result and previous reports, we found that *npr-1* loss of function in an otherwise Bristol background significantly increased the foraging speed of these strains on food (Fig. 5A). Additionally, nearly isogenic line (NIL) animals with the Bristol *npr-1* allele in an otherwise Hawaiian genetic background moved as slow as the Bristol strain, and NIL animals with Hawaiian *npr-1* allele in an otherwise Bristol background moved faster on food, resembling the Hawaiian strain (Fig. 5A). Thus *npr-1* is likely to be the causative gene underlying the observed speed differences. Using the same set of NILs, we also identified *npr-1* as the gene contributing to variation in deceleration (Fig. 5A), consistent with the strong phenotypic correlation of this trait with speed at early time points (Spearman rho=0.9).

**Figure 5:**
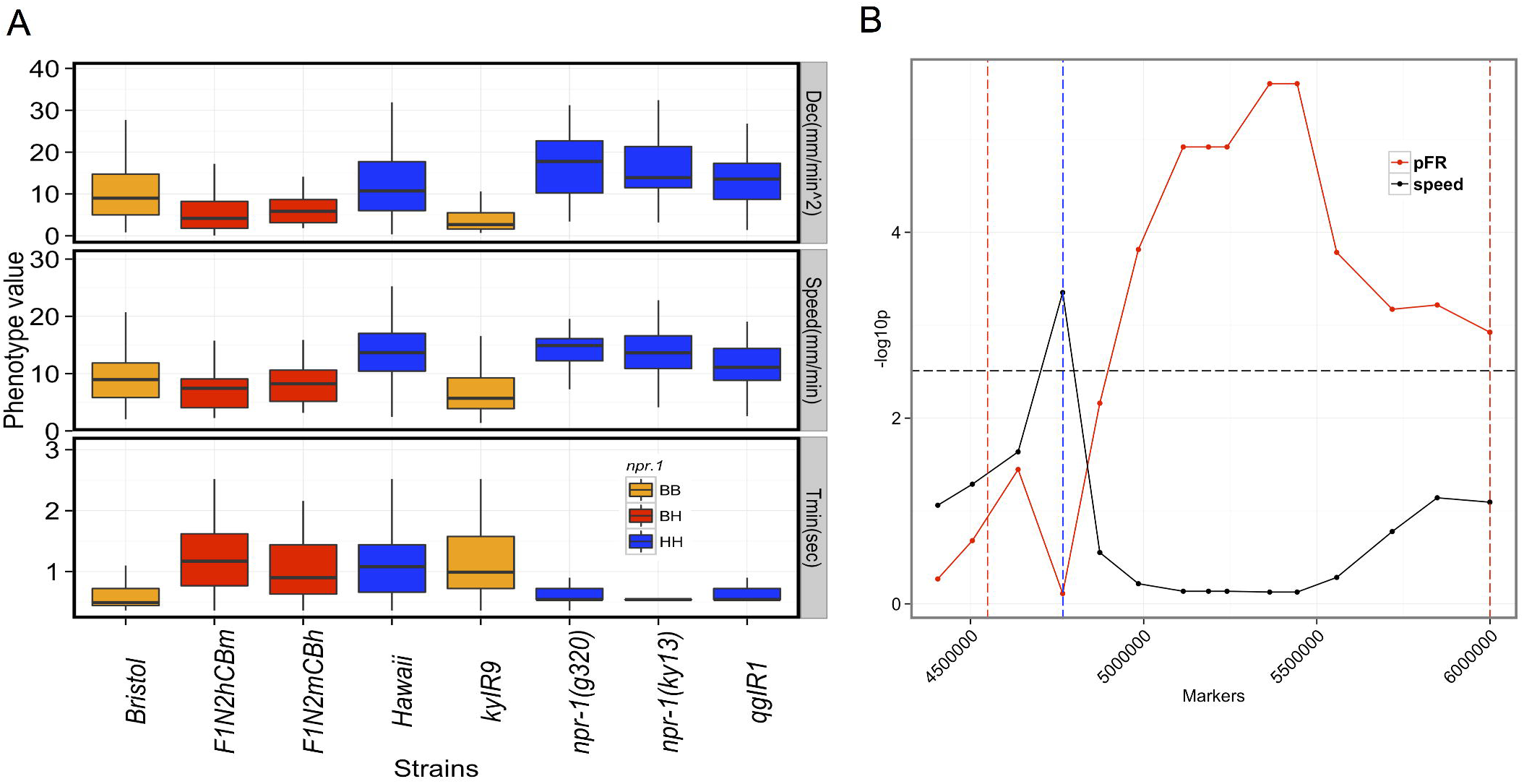
A) Distributions of three components of the avoidance behavior for parental, introgression and *npr-1* loss-of-function strains. Boxplots of deceleration (Dec, upper panel), speed at early time points (speed) and time to pause state (Tmin) for the indicted strains. Briatol and Hawaii are the parental strains. F1N2hCBm: F1 heterozygotes obtained by crossing Bristol hermaphrodites with Hawaiian males. F1N2mCBh: F1 heterozygotes obtained by crossing Bristol males with Hawaiian hermaphrodites. *kyIR9* is an introgression line with Bristol like *npr-1* in an otherwise Hawaiian background. *qgIR1* is an introgression line with Hawaii like *npr-1* in an otherwise Bristol background. The npr-1 status are color coded as follows: orange: Bristol *npr-1* allele; blue: Hawaii like (loss-of-function) *npr-1* allele and BH: F1 heterozygotes from Bristol and Hawaii cross. **B) Speed and probability of responding by reversals are genetically separable.**–log10 of p-values (Y-axis) resulting from t-test at each marker (X-axis) , for the presence of a loci for speed at early time point (speed, black) and probability of responding by reversal (pFR,red) (see methods for more details). Black horizontal line: Bonferroni corrected threshold for fourteen markers (black or red dots). The blue vertical dashed line represents the position of *npr-1*. The red vertical dashed lines represent the 1.5 LOD drop interval of the proximal QTL on X.

We detected a significant QTL for mean time to pause state (Tmin) that overlapped with the QTL for speed at early time points and for deceleration. However, unlike with deceleration and speed, we found no evidence for *npr-1* contributing to Tmin. This result is consistent with a weak correlation of Tmin with speed and deceleration (Spearman’s rho= .22 and -0.05, respectively). Strains with loss of function mutations in *npr-1* or with the Hawaiian form of *npr-1* exhibited Tmin that resembled the Bristol strain, suggesting that this trait is independent of *npr-1* (Fig. 5A). Thus, allelic variation at both *npr-1* and one or more closely linked genes contribute to the observed variation in different aspects of the avoidance response.

Next we focused on fine mapping QTL that contributed to variation in probability of responding by reversals (pFR). Several lines of evidence suggested that the causative gene(s) underlying pFR is unlikely to be *npr-1*. Animals with *npr-1(lf)* mutations or Hawaii like *npr-1* alleles in a Bristol background behaved like the Bristol parent (Table S9). The pFR of a NIL with Hawaii like *npr-1* allele in an otherwise Bristol background was similar to the Bristol parent (Table S9). Finally, the correlation between pFR and the *npr-1*-dependent speed at an early time point in the RIAILs was relatively weak (Spearman rho=-0.32,p=0.0002).

To fine-map the loci underlying pFR, we generated a panel of thirty-one X-chromosome introgression lines (NILs) and genotyped them by whole genome sequencing (Fig S8, See methods). We quantified pFR and speed at the earliest time point of the assay (speed at 0.18s). As with the RIAILs, we found little correlation between these two traits among the introgression lines (Spearman’s rho: 0.18, p=0.32). We examined the effect of *npr-1*, the genetic background, and QTL_Xb on the variation in speed and pFR in the NIL panel using ANOVA. We found that *npr-1* contributed significantly to the variation in speed but not to variation in pFR among these introgression lines (see Methods, Table S10 and S11). Additionally, the genetic background of an introgression line and a distal QTL on chromosome X contributed significantly to variation in pFR but not to speed. Taken together, these data reveal a separate genetic basis for pFR and speed. To further narrow the QTL interval, we next sought to identify markers within the QTL interval that were significant predictors of pFR. Consistent with separate genetic loci for speed and pFR, we found that markers immediately adjacent to *npr-1* were a significant predictor of speed, whereas markers distal to *npr-1* significantly contributed to the variation in pFR (Fig. 5B). Thus, the gene or genes contributing to variation in pFR are distinct from *npr-1* and likely lie within a 750 kb interval distal to *npr-1*. Given the significant effects of genetic background and a second distal QTL on chromosome X, further studies with a larger set of introgression lines are needed to localize the causative gene.

## Discussion

Most studies investigating the genetic basis of animal behavior have focused on a single genetic background—often the reference laboratory strain. As a result, we have relatively little understanding of the genetic basis of natural variation in behavior. In this study, we sought to identify the genetic mechanisms that generate natural variation in behavioral responses to a thermal stimulus. Our results show that the escape behavior induced by thermal stimuli is composed of simpler behavioral components that are influenced by at least six distinct genetic loci. Our results reveal a genetic system that allows independent modification of behavioral components regulating escape response. Overall, our study reveals the constraints and the flexibility of the genetic mechanisms that generate diversity in *C. elegans* escape behavior.

We found that the genetic basis of intraspecific variation in an escape behavior is relatively simple. A few genetic loci, acting largely in an additive fashion, contribute to the observed behavioral diversity. For example, we show that it is possible to decouple the genetic mechanisms underlying acceleration and deceleration in response to thermal pulse stimuli. Allelic variation at specific loci can alter some behavioral components but not others, demonstrating that the genetic architecture of the escape response is modular to a large degree. Our findings are similar to those recently published on burrowing patterns in deer mice and schooling behavior in sticklebacks [19,20], providing initial evidence that modular regulation of variation in behavior may be common. These similarities in genetic architecture are striking given the diverse organisms and behaviors studied. Similar genome-wide studies of integration of characters at the genomic level have been conducted for morphological traits [32,33,36].

Although we provide strong evidence that distinct genetic loci underlie different components of escape behavior in our RIAIL population, we were less successful in identifying such a set of loci through genome-wide association studies in wild isolates of *C. elegans*. This could be attributable to a lack of power in the association studies with the relatively small number of available isolates and/or to rare variants contributing to variation in thermal stimulus induced escape behavior. Further genetic mapping studies with pairs of wild isolates will elucidate the genetic underpinnings of natural diversity in avoidance behavior and provide novel insights into the genetic basis of a complex behavior.

We further investigated the genetic architecture of escape behavior by focusing on two strains, a wild Hawaii-derived strain and the commonly used laboratory N2 strain. The RIAIL population we used was constructed from a Hawaiian wild isolate and the laboratory strain N2. It has become increasingly clear that laboratory-derived polymorphisms in a G-protein coupled receptor gene, *npr-1*, have large effects on the life history and behavior of N2 [26,29,30,34]. *npr-1* has also been implicated in high temperature (ΔT > 9°C) avoidance behavior in *C. elegans* [35]. Consistent with previous reports, we observed that *npr-1* variation underlies speed differences between the Hawaii and Bristol strains. In addition, *npr-1* also contributed to variation in deceleration upon exposure to a thermal pulse stimulus. Studying wild isolates beyond the laboratory strain will provide novel insights masked by the effects of *npr-1*. This is highlighted by our finding of a locus on chromosome X, distal to the *npr-1* locus that contributes to variation in un-stimulated speed in the wild isolates.

Although the QTL associated with time to pause state and probability of responding by reversal overlaps with *npr-1*, our analysis suggests that these are *npr-1*-independent traits. We found little correlation between the *npr-1* genotypes and these traits. Another candidate for decision making behavior on the X chromosome is the tyramine-gated G-protein-coupled receptor *tyra-3*. Polymorphisms in *tyra-3* have been shown to contribute to variation in decision making between Bristol and Hawaii strains in foraging [34]. However, this gene is not located in the fine-mapped interval, and is thus unlikely to regulate escape behavior.

We also detected a QTL on chromosome V contributing to reversal duration and frequency of body bends during reversals. This locus coincides with the location of *glc-1*, a glutamate-gated chloride channel family gene. Loss-of-function mutations in this gene have been shown to impair reversals and also to cause resistance to the anthelmintic abamectin [37]. Consistent with this, we found that reversal duration and frequency of body bends during reversals were significantly correlated with abamectin resistance in the RIAILs (unpublished observations). This implies that *glc-1* may integrate reversal duration, number of body bends during reversal state, and probability of omega turns. Different allelic forms of *glc-1* may result in differential execution of avoidance behavior in response to thermal stimuli.

Our results from RIAILs provide new insights into the genetic mechanisms of escape behavior in *C. elegans*. We found that QTL on chromosome V and X contribute to the duration of reversals (‘timer’), whereas QTL on chromosomes IV and V contribute to variation in the number of body bends during reversals (‘counter’). Hence, changing the causative allele(s) on chromosome X or IV can independently modify the timer or the counter during reversals. Changing the causative allele at the QTL on chromosome V will affect both aspects of the reversal and hence may act as a constraint. We also provide evidence that the probability of responding by reversals and the probability of making an omega turn are influenced by unlinked loci, suggesting that there are independent neuronal routes to altering each of these behavioral components. Ultimately, differences in escape behavior at the genetic level will be reflected at the level of neuronal circuitry and/or neuroanatomy. The present study lays the groundwork for future evolutionary, genetic and neuronal circuitry studies to relate the genetic differences to individual variations in escape behavior.

## Materials and Methods

**Strains:** All strains used in this study were maintained under standard nematode culture conditions. Unless mentioned otherwise all animals assayed were of the hermaphrodite sex. The strains assayed for the calculation of various thermal response behavioral metrics are listed in Data S12.

**Behavioral assays:** Unless otherwise mentioned mid-late L4 stage worms were picked on 6 cm standard NGM plates with agar ~24 hours before the assay. Assay plate preparation: 10 cm plates containing 10 ml of agar medium [ 17 gm of agar (Difco, Detroit, MI), 2.7 gm of Bactopeptone (Difco), 0.55 gm of Tris base (Sigma, St. Louis, MO), 500μl of 1(M) Tris HCl (Sigma), 2.0 gm of NaCl (Fisher Scientific, Pittsburgh, PA), and 1 ml of ethanol containing 5 mg/ml cholesterol (Sigma), per liter H2O]. Plates were stored at 4°C and used within two weeks.

100μl of E.coli (op50) OD ~0.6-0.8 was spread evenly on the agar plates at least 16 hours prior to the assay. On the day of the assay single worms were transferred to the agar plates seeded with bacteria and kept at 20°C for at least 10 minutes. The worms were then subjected to thermal stimulus as described below. We computed a given behavioral metric obtained from 30 (on an average) individual worms per strain. The numbers of worms per strain are depicted in Data S12)

**Thermal stimulus assay:** Worms were imaged using a Leica MZ16APO stereomicroscope and a Basler firewire CMOS camera (A602fm; Basler, Ahrensburg, Germany). A collimated beam with a 1/e diameter of 1.50 mm from a 1440 nm diode laser (FOL1404QQM; Fitel, Peachtree City, GA) was positioned to heat the area covering the worm. The diode laser was driven with a commercial power supply and controller (LDC 210B and TED 200C; Thorlabs, Newton, NJ). A custom program written in LabVIEW (National Instruments, Austin TX) was used to control the firing, power, and duration of the IR laser, while simultaneously recording images of the crawling worm for 15 seconds at 14 or 28 Hz. Images were processed offline using custom programs written in LabVIEW and MATLAB (Mathworks, Natick, MA)[7]. A thermal camera (ICI 7320, Infrared Camera Inc., Texas, USA) was used to measure the temperature of the agar when heated by the IR laser. We were able to generate thermal pulses with a temperature change ranging from ΔT= 0.4°C to 9.1 °C with ramp rates ranging from 0.8°C/sec to 18°C/sec respectively. These ΔT were measured from the baseline temperature (or room temperature), which was typically ~20°C.

Behavioral Quantification: Speed profile of each worm over time was generated by a custom written program in MATLAB and LabVIEW that monitored the position of the Center of Mass (COM) of the worm image [7]. From the position of COM in the sequence of captured images, the speed-related traits namely the speed at every time point, acceleration and deceleration of the worm were identified using custom code. All analysis was performed separately on the 14 frames per second (fps) and 28 fps data. Once processing was complete, the 14 fps data was converted from frames to time (in seconds) and combined with the 28 fps data to produce the final results. Some of the traits namely the reversal duration, number of body bends during reversals and probability of responding by reversals and omega turns were quantified manually from the movies of individual worms. A thermal pulse was applied to a forward moving worm and any movement opposite to the direction of the movement of the worm after application of the thermal pulse was considered a reversal. The number of body bends and omega turns were assigned following previously published criteria [38].

### Genomic DNA library preparation and sequencing

Genomic DNA was prepared using the DNeasy Blood and Tissue Kit (Qiagen). DNA concentration was determined using the Broad-range Quant-it dsDNA High-Sensitivity DNA quantification kit (Invitrogen). DNA was diluted to 1.7 ng/μl. Libraries were prepared by the Nextera tagmentation procedure using the Nextera HMWbuffer according to manufacturer’s (Epicentre) protocol. The transposition reaction was performed for 5 min at 55°C and the resulting samples were purified using the MinElute kit (Qiagen). Fifteen microlitres of the purified fragmented DNA was PCR-amplified and barcoded with custom 5-base pair (bp) sequences using Ex Taq polymerase (Takara) for 10 PCR cycles. Five μl of each PCR-amplified sample was pooled into one 8 or 16 plex library. The pooled library was loaded on a 2%agarose gel, and the 350–550 bp region was excised and gel-extracted using QIAquick Gel Extraction Kit (Qiagen). Final libraries were diluted to 3.3 ng/μl and sequenced using the paired-end module on a HiSeq 2000 (Illumina).

**NIL construction and genotyping:**

The sources of the (Nearly Isogenic Lines) NILs and the methods use to generate them are outlined in the Data S12. For genotyping, we used custom R and python code to demultiplex and trim the ends of the barcoded sequencing data. Sequencing reads were assigned to each strain based on the 5-bp barcode at the beginning of each read. The internal 19-bp transposon sequence and 10 bp on the right end of each read were removed. Reads were aligned to the WS210 elegant reference genome using the Burrows–Wheeler Aligner using default parameters. Next SAMtools/BCFtools was run to generate pileup and merged vcf files of the NILs for the strains using default parameters. Genotype likelihoods for the N2 and CB4856 alleles for each genotypic variant were extracted from the VCF file as follows.

We ran vcftools to extract variant sites known to be different between N2 and CB4856. We extracted the genotype likelihoods for the homozygous reference and alternate calls. Then we used a hidden Markov model and the Baum-Welch algorithm to calculate the posterior likelihood that a variant is coming from N2 or CB4856. Variants with a higher likelihood of coming from N2 relative to coming from CB4856 were coded as N2 and vice versa. We obtained genotype information of 16853 positions on chromosome X as either belonging to N2 or CB4856 and deduced the breakpoints from the resulting data. We also independently identified breakpoints on chromosome X for the NILs using multiplex shotgun genotyping (MSG) with concordant results. For each strain, two million reads were randomly subsampled to improve the speed of the computation pipeline, which requires very low per base coverage (Andolfatto et al. 2011). Briefly, reads were mapped to both parental genomes and only ancestry informative markers were used to impute ancestry along the chromosome. Because lines vary in their proportions of each parent on the X chromosome, naïve priors were used (prior parent 1: 0.5, prior parent 2: 0.5). The raw output of the MSG pipeline is posterior probabilities for each genotype (parent1, heterozygous, parent2), and information was obtained at 75430 markers that were used to infer the breakpoints.

**Statistical analyses:**

QTL mapping: We used 138 advanced intercross recombinant inbred lines (RIAILs) made from N2 and CB4856 strains as described elsewhere for QTL mapping. QTL mapping was performed using R/QTL. Linkage of the escape behavior components (measured by procedures described above) to 1454 markers across the genome was determined under a single-QTL model in an interval-mapping framework. Genome-wide LOD thresholds for mapping a given trait was determined using 1000 random permutations of the data for the RIAILs for the trait in question. 1.5 LOD support intervals were used to determine the confidence intervals for each QTL. To account for large effect QTL, markers corresponding to these were included as covariates and single-QTL analysis performed again. For multiple QTL mapping of normally distributed traits, we looked for any significant epistatic as well as additive interactions with the markers corresponding to the maximum LOD scores in the chromosomes where the QTL were detected for a given trait in R/QTL. For example, for a trait with two QTL, the model formula was y~ Q1+Q2+Q1:Q2, where Q1 and Q2 represent the QTL objects corresponding to the markers at the maximum LOD scores of the significant QTL peaks. Q1:Q2 is the interaction term between these two QTL objects. We also performed two-dimensional scans as implemented in R/QTL but failed to detect any epistatic interactions. These fit of these models were used to estimate the contribution of the QTL to the total phenotypic variation in the RIAIL population. We partitioned total phenotypic variation into genotypic variation (VG; i.e. among RIAIL or Wild isolate) and error variation(VE), using a random-effects analysis of variance and the broad sense heritability was estimated for each trait separately for the RIAIL or the wild isolate populations [39].

**GWAS:** Association mapping was carried out with 4690 SNPs across the genome using EMMA [40] as described in more detail elsewhere [14].

**Fine mapping using nearly isogenic lines:** To phenotype escape behavior, we performed ~ 30 replicates per nearly isogenic line to determine the mean speed at 0.18s(sp0.18) and the probability of responding by a reversal(pFR). To determine if background of the NILs or the *npr-1 locus* has any effect on the speed at 0.18s and pFR traits we performed an analysis of variance using the formula trait~background+npr1+background*npr1 to identify significant contributors for a given trait. The results of this analysis are shown in Table S10 and S11. Next, to narrow the interval encompassing the identified QTL on chromosome X we tested for the association between 15 markers on the X chromosome (spanning intervals ~4405000 to ~6000000 bp) and the traits (sp0.18s and pFR) using a linear model taking into account the effect of the genetic background and the distant QTL identified on X (QTL_Xb). The strength of association (-log10p) resulting from the fit for each marker is depicted in Fig. 5. The significance threshold was determined by Bonferroni correction −log(0.05/15).

## Acknowledgements

The authors are grateful to Frank Albert, Erik Andersen, Alejandro Burga, Meru Sadhu, Sebastian Treusch, Xin Wang and Danny Zeevi for helpful discussions and comments on the manuscript. We also thank Andres Bendesky and Matthew Rockman along with the *Caenorhabditis* Genetics Center, which is funded by the NIH National Center for Research Resources (NCRR), for some nematode strains used in this work. Some strains were provided by the CGC, which is funded by NIH Office of Research Infrastructure Programs (P40 OD010440).

## Supplementary materials

**Figure S5: A) Spread of** heritabilities of the sixty-six behavioral components in the wild isolate and the RIAIL population in response to ΔT~0.4°C. **B)** Correlation of six components of the avoidance behavior among wild isolates. The behavioral components are: sp0.18s: speed at 0.18 seconds, sp1.61s: speed at 1.61 seconds, sp8.1s: speed at 8.1 seconds after start of the assay, pFR: probability of responding by reversals, W=number of body bends during reversals, pRO: probability of inducing omega turns. The magnitude of significant Pearson correlation coefficients (p< 0.01) among these behavioral metrics are shown for wild isolates in circles. The phenotypes are hierarchically clustered and cut into two clusters bordered by red lines. **C,D)** Common naturally occurring variation on chromosomes IV and X contribute to natural variation in speed at early time points (C) and acceleration (D) during escape behavior. Plots of the results of genome wide association mapping for acceleration and speed at 0.2 seconds. Significantly associated markers are indicated by red lines.

**Figure S6: Linkage analysis of various escape behavioral components related to centroid speed.** Logarithm of the odds (LOD) scores plotted against marker positions. Each facet represents a chromosome depicted as I through V and X. The traits that are mapped are shown on the title of the corresponding figure. The horizontal dashed line represents the 5% genomewide significance threshold obtained after 1000 permutations. A-D) LOD profiles for centroid speed at particular time points. E and F) LOD profiles for centroid speed at every 0.18 seconds for chromosomes IV (E) and X(F) respectively. Each facet represents the mapping results of speed at a single time point (depicted as sp*.**s in red). The red horizontal line represents the 5% genomewide significance thresholds. The gray vertical line in (E) represent the peak marker position of the significant QTL. The blue vertical line in (F) represents the position of *npr-1* gene. Asterisks represent the speed at time points where a significant QTL was detected on chromosome IV. G-J) QTL analysis results of the speed related escape behavior components. The axes of the plots are same as in Fig S6A. The traits mapped are in the title of each plot.

**Figure S7: Linkage analysis of various escape behavioral components related to reversals and omega turns.** The axes of the plots are same as in Fig S6A. The traits mapped are in the title of each plot (A-E).

**Figure S8: A)** Overlay of the speed (black) and the probability of responding by reversals (red) QTL on chromosome X. The horizontal dashed lines represent genomewide significance thresholds. The vertical dashed lines represent the peak of the respective QTL.

**B)** The breakpoints on the X-chromosomes (X-axis:physical position in basepairs) of introgression lines (Y-axis) as determined by whole genome sequencing. Blue indicates Hawaii markers whereas orange indicates Bristol markers. The vertical red-dashed line represents the position of *npr-1*. The black and the white vertical dashed lines represent the proximal and distal QTL intervals respectively.

## Tables

**Table S9: *npr-1* does not contribute to probability or responding by reversals**

**Table S10: ANOVA results: Effects of genetic background, *npr-1* and distal QTL on chromosome X (QTL_Xb) on speed**

**Table S11: ANOVA results: Effects of genetic background, *npr-1* and distal QTL on chromosome X (QTL_Xb) on probability of responding by reversals**

## Movies

**Video S1**: Avoidance behavior of Bristol animal in response to ΔT~0.4°C

**Video S2:** Avoidance behavior of Hawaiian animal in response to ΔT~0.4°C

**Video S3:** Avoidance behavior of Bristol animal in response to ΔT~9.1°C

**Video S4:** Avoidance behavior of Hawaiian animal in response to ΔT~9.1°C

## Data

**Data S12.xlsx: Genotype and phenotype details for RIAILs, Wild isolates and NILs.**

